# Evidence for a ‘trap-and-flip’ mechanism in a proton-dependent lipid transporter

**DOI:** 10.1101/2021.08.05.453402

**Authors:** Elisabeth Lambert, Ahmad Reza Mehdipour, Alexander Schmidt, Gerhard Hummer, Camilo Perez

## Abstract

Transport of lipids across membranes is fundamental for diverse biological pathways in cells. Multiple ion-coupled transporters participate in lipid translocation, but their mechanisms remain largely unknown. Major facilitator superfamily (MFS) lipid transporters play central roles in cell wall synthesis, brain development and function, lipids recycling, and cell signaling. Recent structures of MFS lipid transporters revealed overlapping architectural features pointing towards a common mechanism. Here we used cysteine disulfide trapping, molecular dynamics simulations, mutagenesis analysis, and transport assays *in vitro* and *in vivo*, to investigate the mechanism of LtaA, a proton-dependent MFS lipid transporter essential for lipoteichoic acids synthesis in the pathogen *Staphylococcus aureus*. We reveal that LtaA displays asymmetric lateral openings with distinct functional relevance and that cycling through outward- and inward-facing conformations is essential for transport activity. We demonstrate that while the entire amphipathic central cavity of LtaA contributes to lipid binding, its hydrophilic pocket dictates substrate specificity. We propose that LtaA catalyzes lipid translocation by a ‘trap-and-flip’ mechanism that might be shared among MFS lipid transporters.

---

Major facilitator superfamily (MFS) transporters are found in all kingdoms of life and move a large variety of molecules across biological membranes [1–8]. Structural characterization of MFS transporters that participate in the uptake of water-soluble molecules and extrusion of drugs has contributed to a broad understanding of their transport mechanism [4, 8–17]. However, multiple reports have attributed alternative functions to MFS transporters, such as translocation of lipids associated with fundamental biological pathways. Some examples include the bacterial lysophospholipid transporter LplT, involved in lipids recycling in Gram-negative bacteria [7, 18]; the human transporter MFSD2A, expressed at the blood-brain- and blood-retinal-barrier, contributing to major uptake of docosahexaenoic acid (DHA) [5, 6, 19–21]; the human transporters Spns2 [22, 23], and MFSD2B [24], which contribute to transport of sphingosine 1-phosphate (S1P) in endothelial cells and erythrocytes; and the gentiobiosyl-diacylglycerol transporter LtaA, involved in cell wall synthesis in *Staphylococcus aureus* [25, 26]. However, despite their well described cellular roles, the mechanisms of MFS lipid transporters remain insufficiently understood.

We have previously shown that LtaA is a proton-dependent MFS lipid antiporter [26]. It contributes to adaptation of *S. aureus* to acidic conditions, common in the skin and nasopharynx of the human host [26–28]. LtaA takes part in the assembly of lipoteichoic acids, phosphate-rich polymers important for control of bacterial cell division, protection from environmental stress, host cell adhesion, antibiotic resistance, biofilm formation, and immune evasion [29–32]. *S. aureus* lipoteichoic acids display a polymer of 1,3-glycerol-phosphate repeat units attached to C-6 of the non-reducing glucosyl of the glycolipid gentiobiosyl-diacylglycerol [31–33]. This glycolipid is synthesized at the cytoplasmic leaflet of the membrane by the glycosyltransferase YpfP, and is translocated to the outer leaflet by the activity of LtaA [25, 26]. The essential role of LtaA in adjusting the pool of glycolipids available at the extracellular side of the membrane, makes this protein a central player for lipoteichoic acids assembly and a potential target for drugs aiming to counteract antimicrobial resistant *S. aureus* strains e.g., methicillin-resistance *S. aureus* (MRSA) and vancomycin-resistant *S. aureus* (VRSA) [30].

Two different general models of transporter-catalyzed lipid translocation have been proposed in the past [34–42]. A ‘trap-and-flip’ model, in which the lipid substrate is retrieved from one leaflet of the membrane, enclosed into a central cavity, and then delivered to the other leaflet [40, 43, 44], and a ‘credit-card’ model that departs from the classical alternating-access model and involves translocation of the lipid head-group across a hydrophilic cleft or cavity in the transport protein, while aliphatic chains remain embedded in the membrane [36–38, 41, 42, 45, 46]. However, it is not known which of these two models describe better the mechanism of MFS lipid transporters. Answering this question is not only important to understand the basis of the processes catalyzed by these proteins, but could also provide a foundation for the design of drugs and/or lipid-linked-bioactive molecules targeting cells or organs expressing pharmacologically relevant proteins from this superfamily.

Until now, only the high-resolution structures of outward-facing LtaA and inward-facing MFSD2A have been elucidated [21, 26]. Both transporters displayed the canonical MFS fold of 12 transmembrane (TM) helices and an amphipathic central cavity that has not been observed in any MFS transporter of water-soluble molecules. The similar architectural features observed in the structures of LtaA and MFSD2A indicate common elements in their transport mechanisms and likely among all MFS lipid transporters. Here, we used cysteine disulfide trapping of outward- and inward-facing LtaA, in combination with molecular dynamics simulations, mutagenesis analysis, and transport assays *in vitro* and *in vivo*, and showed that cycling through outward- and inward-facing conformations is essential for LtaA activity. We demonstrate that LtaA displays membrane exposed lateral openings with distinct functional relevance and characterized the architecture and biochemical properties of the amphipathic central cavity during alternating-access. Our results indicate that while the hydrophilic pocket of the amphipathic central cavity dictates substrate specificity, the hydrophobic pocket is only relevant for aliphatic chains binding. We describe critical mechanistic elements revealing that LtaA adopts a ‘trap-and-flip’ mechanism that might be shared among MFS lipid transporters.

## Results

### Model of inward-facing LtaA and validation by cysteine cross-linking

To investigate whether LtaA uses a ‘trap-and-flip’ or a ‘credit-card’ mechanism, we first aimed to establish a system that allowed us to perform cysteine disulfide trapping of end-point conformations of LtaA during its transport cycle. The architecture of the previously solved structure of LtaA [26], facilitates cysteine disulfide trapping of outward-facing states, whereas there is no structural information to guide trapping of inward-facing states. Thus, we first generated an inward-facing model of LtaA using ‘repeat-swap’ modeling [47]. Like other transporters from the MFS superfamily, the topology of LtaA consists of two domains, a N-terminal domain (TM1-TM6; domain-1), and a C-terminal domain (TM7-TM12; domain-2), each of which contains two structural repeats with inverted-topology related by a pseudo-rotational two-fold symmetry axis parallel to the plane of the membrane (**Fig. 1A,B**). After swapping the conformations of the inverted repeats observed in the outward-facing structure of LtaA (PDB ID 6S7V) [26, 47], we constructed a large set of models *in silico* that were refined aiming to improve side chains packing, stereochemistry, and modeling scores. The models with the best scores converged to one conformation (**Fig. 1C and Suppl. Table. 1**), which displayed multiple interactions between the extracellular parts of TM1-TM7, TM2-TM11, and TM5-TM8, sealing the entrance to the central cavity (**Fig. 1C**). In contrast, the cytoplasmic regions of helices TM2-TM11, TM5-TM8, and TM4-TM10, lining the entrance to the central cavity from the cytoplasm, are away from each other about 16.0±0.1 Å, 16.0±0.1 Å, and 17.6±0.2 Å, respectively (**Fig. 1C**). The helical loop between TM6 and TM7 that connects the N- and C-terminal domains was modeled based on the conformation observed in the outward-facing structure.

**Figure 1.**
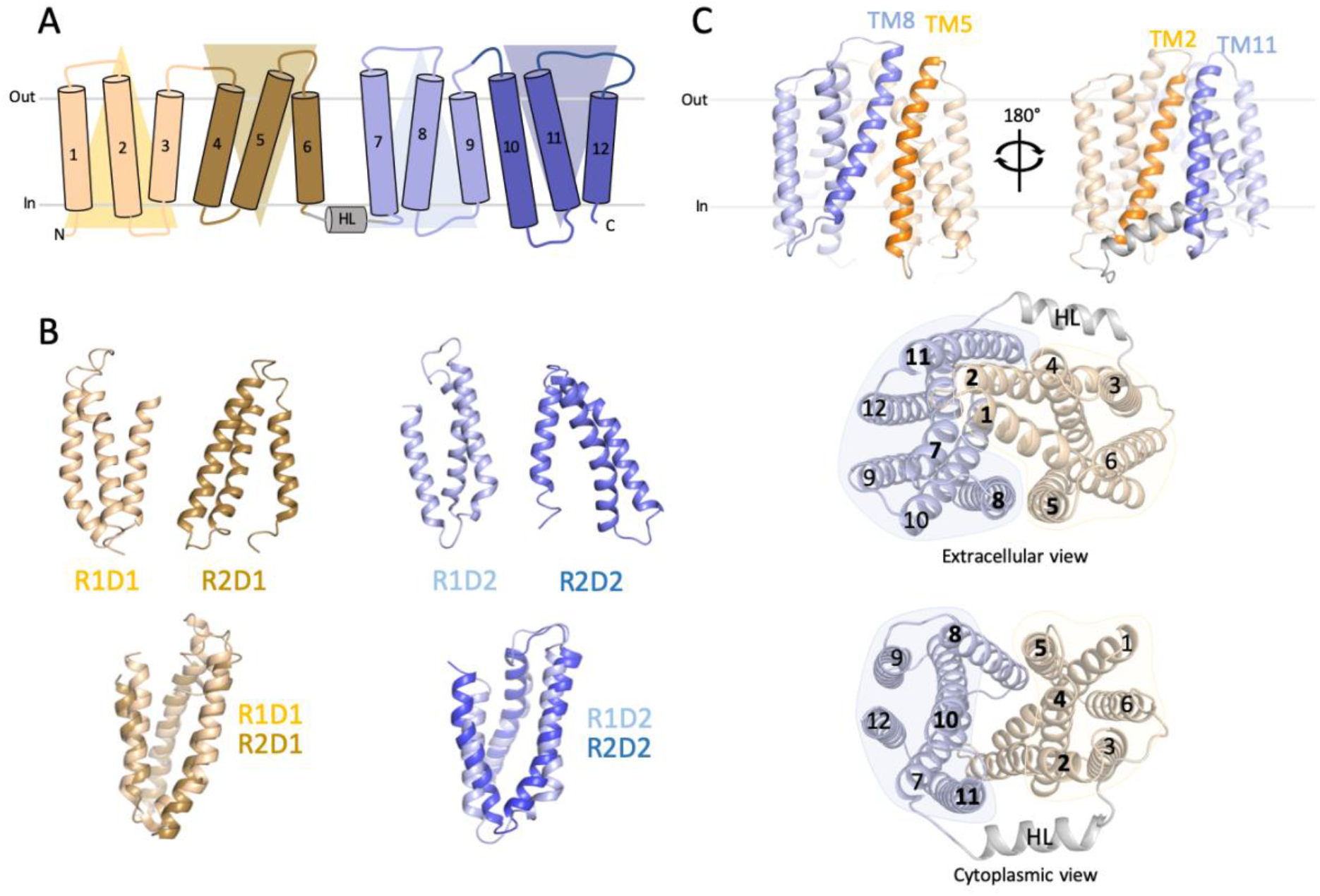
‘Repeat swap’ modeling of inward-facing LtaA. **A**. Topology representation of LtaA. Domain-1 (N-terminal) and domain-2 (C-terminal) are shown in light orange and light blue, respectively. **B**. Individual repeat domains as observed in outward-facing LtaA (PDB ID 6S7V), and superposition of inverted repeats (r.m.s.d. = 2.5 Å and 3.0 Å for aligned Cα atoms of R1D1/R2D1 and R1D2/R2D2, respectively). R1D1 and R2D1 indicate the first and second repeats in the N-terminal domain, respectively, whereas R1D2 and R2D2 indicate the first and second repeat in the C-terminal domain, respectively. Colors are according to panel **A**. **C**. Side-views of inward-facing LtaA model showing TM helices that line the lateral openings. Extracellular and cytoplasmic views are also shown.

To validate the inward-facing model, we selected pairs of residues among the extracellular regions for which Cβ-Cβ distances were less than 7 Å, but which present Cβ-Cβ distances of over 12 Å in the outward-facing structure (**Suppl. Fig. 1**). Based on these criteria, we identified the pairs F45-T253, A53-T366, and K166-I250 (**Fig. 2A and Suppl. Fig. 1**), occupying three different positions that provide good coverage of the conformational change predicted by our models. We then introduced cysteine residues at these positions on a starting construct in which the one native cysteine in LtaA was replaced with serine. The cysteine-less LtaA variant effectively performed glycolipid flipping in proteoliposomes (**Suppl. Fig. 2**). The three mutants F45C-T253C, A53C-T366C, and K166C-I250C were then irreversibly crosslinked with N,N’-(o-phenylene)-dimaleimide (o-PDM), which has a spacer arm length of 6 Å. Crosslinked and non-crosslinked LtaA mutants were digested with either trypsin or chymotrypsin, and analyzed by high-resolution liquid chromatography–mass spectrometry (LC–MS) to evaluate the presence of non-crosslinked cysteine containing peptides. The peptides abundance was normalized against an internal reference peptide. We successfully identified non-crosslinked peptides in untreated samples of the three mutants F45C-T253C, A53C-T366C, and K166C-I250C (**Fig. 2A and Suppl. Fig. 3**). The abundance of these peptides was clearly diminished in the crosslinked protein samples (**Fig. 2A**), demonstrating that the selected pairs of residues are in close proximity as predicted in the inward-facing model.

**Figure 2.**
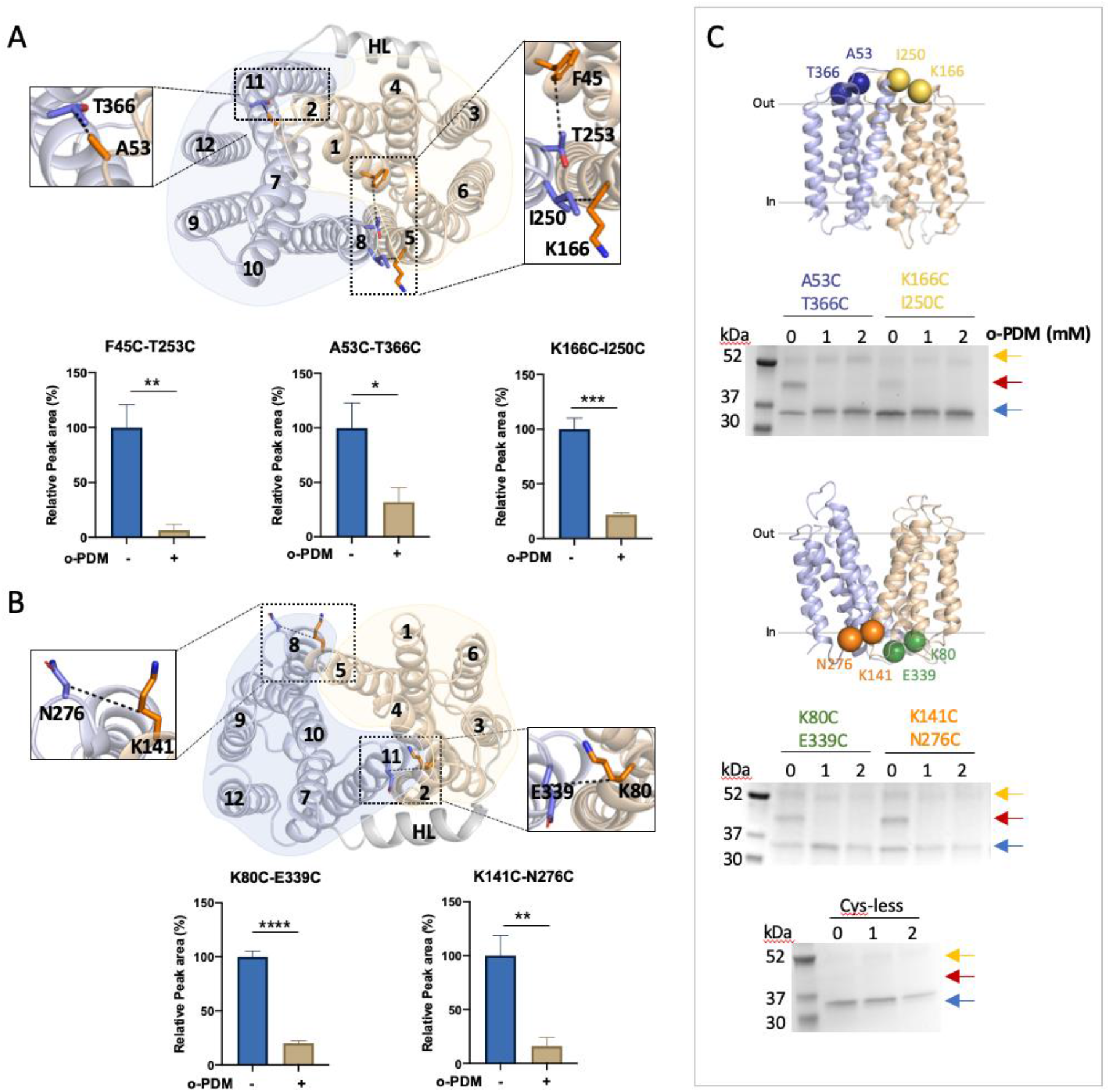
LtaA adopts inward- and outward-facing states. Selected residues for cross-linking of LtaA in inward-facing conformation (**A**) and outward-facing conformation (**B**). N-terminal and C-terminal domains are shown in light orange and light blue, respectively. The relative abundance of cysteine containing peptides in absence (-) or presence (+) of o-PDM is shown in histograms. Collision-induced dissociation (CID) spectrum of cysteine containing peptides and elution profiles of peptide fragments are shown in supplementary figure 3. Error bars indicate standard deviation (s.d.), n≥3. *: P≤0.05, **: P≤0.01, ***: P≤0.001, ****: P≤0.0001. **C**. Cross-linking analysis of LtaA in proteoliposomes. Positions of selected cysteine pairs at the extracellular and cytoplasmic regions of LtaA are shown as spheres. SDS-PAGE show band shifts of samples treated with mPEG5K after irreversible cross-link with o-PDM. Separated species are indicated with arrows. Blue arrow: non-PEGylated LtaA; red arrow: PEGylated LtaA; orange arrow: LtaA dimer.

As a control, we performed a similar experiment but with pairs of residues that were shown to interact at the cytoplasmic region of the outward-facing structure (**Fig. 2B**). Thus, we introduced cysteine residues at the positions K80-E339 and K141-N276, present at the cytoplasmic ends of TM2-TM11 and TM5-TM8, respectively. Cβ-Cβ distances between these residues are smaller than 7.5 Å in the outward-facing structure, but larger than 12 Å in the inward-facing models (**Suppl. Fig. 1**). LC-MS analysis of the double mutants K80C-E339C and K141C-N276C confirmed the proximity of these residues as non-crosslinked peptides are more abundant in untreated samples, whereas in the presence of the cross-linking agent their abundances decrease substantially (**Fig. 2B**). In summary, our cross-linking analysis support the predicted conformation and interactions reported by the inward-facing model of LtaA and indicate the position of residues to guide cysteine disulfide trapping of LtaA conformations.

### Alternating conformations in proteoliposome membranes

We investigated the conformations displayed by LtaA in membranes by evaluating the cross-linking of double cysteine mutants reconstituted in proteoliposomes (**Fig. 2C**). The cysteine pairs reported on the conformation of the TM helices that line the lateral openings, TM2-TM11 and TM5-TM8 (**Fig. 1C**). We screened for successful cross-links by using a gel-shift assay in which we first incubated with the o-PDM cross-linker, followed by treating the proteoliposomes with 5-kDa PEG-maleimide (mPEG5k) [48]. This treatment generates a substantial shift in the protein mobility in polyacrylamide gel electrophoresis as mPEG5k irreversibly binds free cysteines. However, if the introduced cysteines are cross-linked by o-PDM, then they will not react with mPEG5k and no shift in gel mobility would be observed. We evaluated the cross-linking of residues A53C-T366C (TM2-TM11) and K166C-I250C (TM5-TM8), positioned at the extracellular region, and K80C-E339C (TM2-TM11) and K141C-N276C (TM5-TM8), located at the cytoplasmic region (**Fig. 2C and Suppl. Fig. 4A,B**). Before cross-linking, each double-cysteine mutant showed a gel shift after incubation with mPEG5K, thus demonstrating PEGylation of free cysteines (**Fig. 2C and Suppl. Fig. 4C**), albeit the shift of K166C-I250C is less prominent (**Fig. 2C**). In contrast, after treatment with o-PDM, all the double cysteine mutants were protected from PEGylation, thus showing that all mutants were successfully cross-linked. The cysteine-less control LtaA, did not show a gel shift in any of the conditions (**Fig. 2C and Suppl. Fig. 4C**), demonstrating that the shifts observed for the mutants were due to PEGylation of cysteines. These results support that when embedded in the membrane of proteoliposomes, LtaA can adopt conformations where residues at the lateral openings lined by TM2-TM11 and TM5-TM8 display similar distances to those reported by the outward-facing structure and the inward-facing model.

### Alternating-access to the central cavity is essential for function

The cross-linking results described above showed that LtaA can cycle through outward- and inward-facing conformations. However, it is not known whether this is required for function. This is important because some flippases and scramblases use a ‘credit card’ mode of transport, where exposing a side cleft or a cavity to one side of the membrane is sufficient for catalysis of lipid transport across the membrane [36–38, 41, 49]. To answer whether alternating between inward- and outward-facing conformations is important for LtaA activity, we performed copper chloride catalyzed cross-linking of residues located at the lateral openings lined by TM2-TM11 and TM5-TM8, and determined proton-coupled glycolipid transport activity of cross-linked LtaA variants in proteoliposomes (**Fig. 3**). In this assay, the addition of the K^+^-selective ionophore valinomycin generates a membrane potential of about −60 mV, which drives proton influx. Acidification of the lumen of proteoliposomes quenches the fluorophore 9-amino-6-chloro-2-methoxyacridine (ACMA) causing a decrease in the fluorescence [26]. The double-cysteine mutants A53C-T366C (TM2-TM11) and K166C-I250C (TM5-TM8) close the extracellular side openings, whereas the mutants K80C-E339C (TM2-TM11) and K141C-N276C (TM5-TM8) close the cytoplasmic openings (**Fig. 3A and Suppl. Fig. 4A,B**). Our results show that except for K141C-N276C (TM5-TM8), independent cross-linking of the remaining three lateral openings decreases LtaA activity relative to non-cross-linked mutants ( **Fig. 3A**).

**Figure 3.**
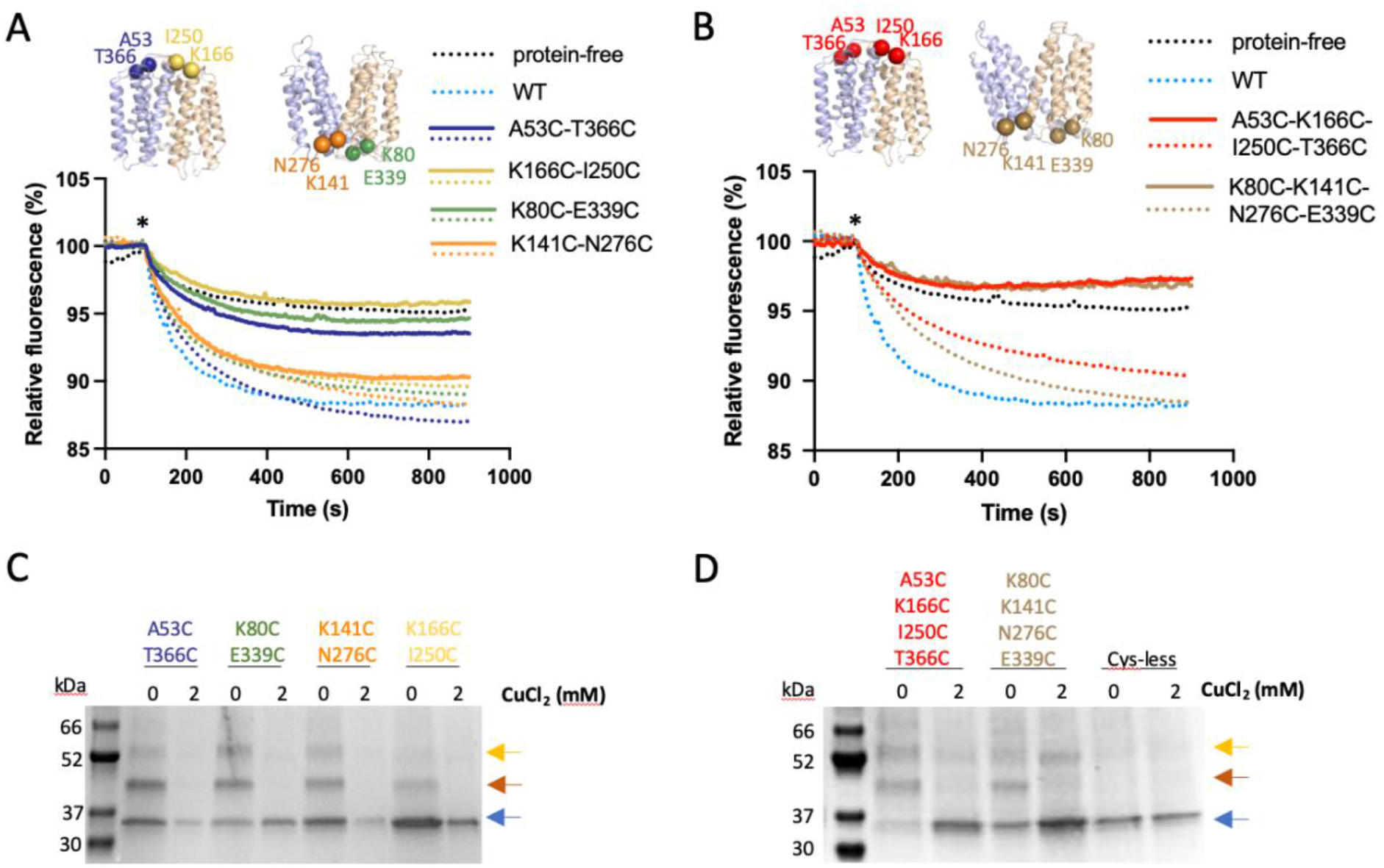
Cycling through outward- and inward-facing conformations is essential for LtaA activity. **A**,**B**. Proton-transport activity of LtaA and variants after chemical crosslinking with CuCl_2_ (solid lines) or in absence of cross-linking treatment (dotted lines) (n≥3). Proteoliposomes and protein-free liposomes containing 100 mM KCl were diluted in buffer containing 10 mM KCl, 90 mM NaCl and ACMA. H^+^ influx was initiated by establishing a membrane potential upon addition of the potassium ionophore valinomycin (asterisk). **C.D**. SDS-PAGE shows band shifts of samples treated with mPEG5K after cross-link with CuCl_2_. Separated species are indicated with arrows. Blue arrow: non-PEGylated LtaA; red arrow: PEGylated LtaA; orange arrow: LtaA dimer.

In addition, we aimed to completely close the cytoplasmic or extracellular cavities and test the effect on LtaA activity (**Fig. 3B**). To do this, we constructed the mutant A53C-T366C-K166C-I250C that after cross-linking would close the extracellular pathway, while the mutant K80C-E339C-K141C-N276C would close the cytoplasmic pathway (**Fig. 3B and Suppl. Fig. 4A,B**). Our results show that in contrast to non-cross-linked proteins, both mutants display background quenching levels, similar to that observed for protein-free liposomes, thus, indicating strong inhibition of transport activity (**Fig. 3B**). We confirmed cross-linking of each double- and tetra-cysteine mutant reconstituted in proteoliposomes by gel shift assays after incubation with mPEG5K (**Fig. 3C,D and Suppl. Fig. 4D**), which showed that after treatment with copper chloride, all the mutants were protected from PEGylation, whereas before cross-linking a gel shift was observed. This confirmed that all mutants were successfully cross-linked in the proteoliposomes samples used in the assay.

Taking together, these results reveal that alternating opening to both sides of the membrane is a requirement for LtaA function. However, not all lateral openings seem to have the same functional relevance. In particular, our results demonstrate that while both extracellular lateral openings are similarly important for function, the cytoplasmic lateral opening lined by TM2 and TM11 has a more significant role, as revealed by the low activity of the cross-linked variant K80C-E339C. In contrast, cross-linking of the cytoplasmic lateral opening lined by TM5 and TM8, does not seem to affect LtaA function strongly.

### Dynamics of lateral openings

For lipid transporters that adopt a ‘trap-and-flip’ mechanism, substrate binding and release involve movement of lipids through lateral openings of the translocation channel [18, 21, 40, 43, 44]. We studied the dynamic behavior of the lateral openings in LtaA when the protein is embedded in a lipid bilayer. To do this, we performed molecular dynamics (MD) simulations of outward- and inward-facing LtaA in a membrane composed of POPG (65%), diacylglycerol (20 %), cardiolipin (10 %), and gentiobiosyl-diacylglycerol (5 %), resembling the membrane of *S. aureus* [50]. During the simulations both states were found to be stable as judged by RMSD plots (**Suppl. Fig. 5**). The simulations revealed that all the optimized inward-facing models exhibit a cavity which is open to the cytoplasm and closed to the extracellular space, whereas the cavity of the outward-facing state is open to the extracellular space and closed to the cytoplasm (**Fig. 4A**). In agreement with the observed wider opening of the extracellular lateral openings, the simulations of outward-facing LtaA showed the intrusion of glycolipid and POPG molecules into the putative translocation pathway (**Fig. 4C**). The glycolipid was seen to intrude from the TM5-TM8 opening, with one of the aliphatic tails reaching to the C-terminal hydrophobic pocket, whereas two POPG molecules intrude from the TM2-TM11 opening (**Fig. 4C**). By contrast, simulations of inward-facing LtaA did not show lipids entering the cytoplasmic cavity.

**Figure 4.**
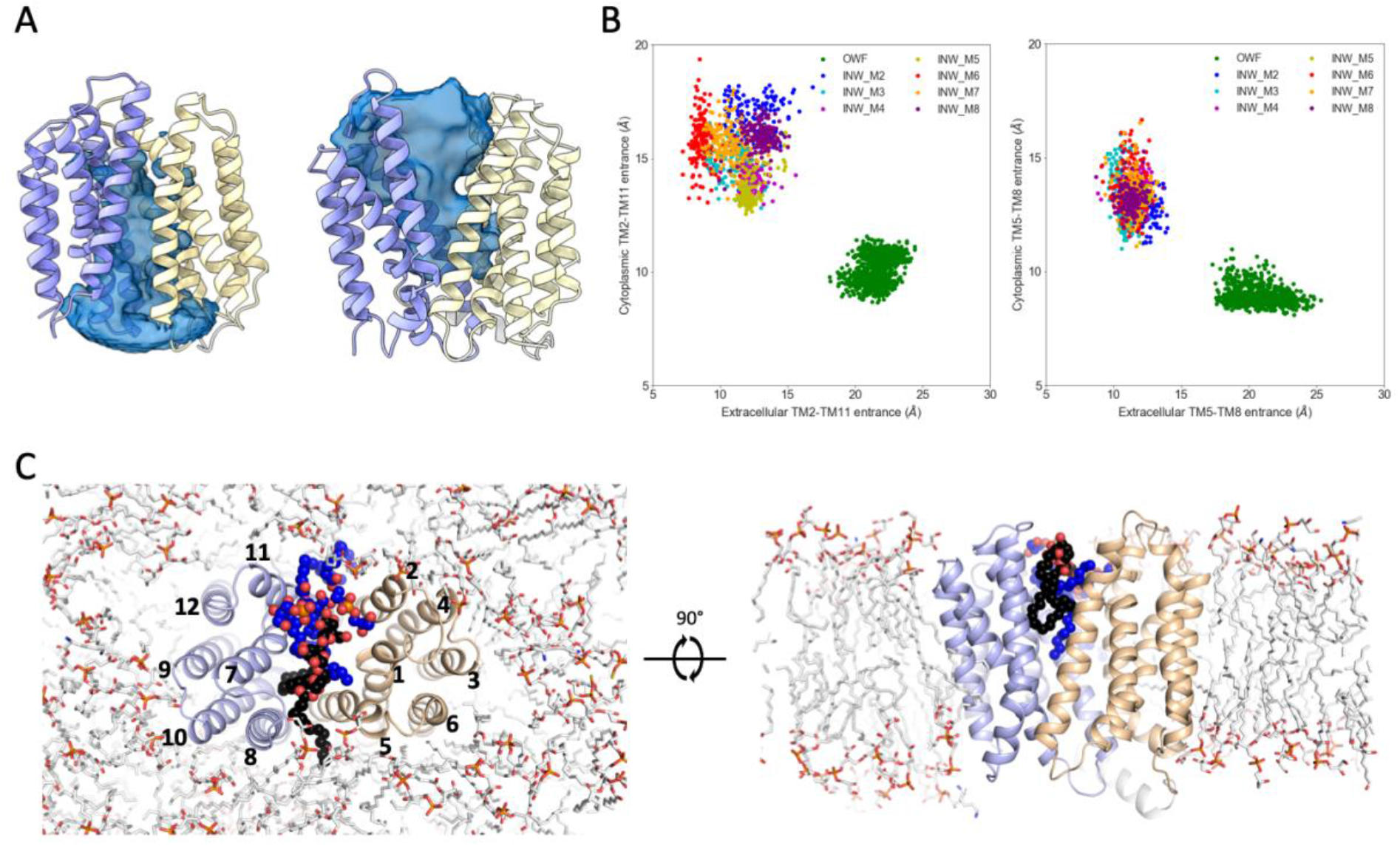
LtaA displays asymmetric opening of cavities and lateral ‘gates’. **A**. Representative view of solvent exposed cavity of inward-facing and outward-facing LtaA as observed during MD simulations. **B**. Analysis of distances between TM helices lining the cytoplasmic and extracellular lateral openings of outward-facing and inward-facing models. The center of masses of the Cα atoms of extracellular residues 52-57 (TM2), 163-167 (TM5), 250-255 (TM8), 364-367 (TM11), and of cytoplasmic residues 77-81 (TM2), 139-143 (TM5), 273-276 (TM8), 341-344 (TM11), were used for the calculation of inter-TM distances. **C**. Intrusion of gentiobiosyl-diacylglycerol (black spheres) and POPG (blue spheres) molecules in the extracellular cavity of LtaA during simulations. N-terminal and C-terminal domains are shown in light orange and light blue, respectively.

In the MD simulations of the inward-facing conformation, we found that the cytoplasmic opening between TM2 and TM11 is wider and more dynamic than that between TM5 and TM8 (**Fig. 4B**). The lateral opening between TM2 and TM11 is thus a likely route for entry of glycolipids into the substrate binding site of the transporter. By contrast, in the outward-facing conformation, the two gates have similar widths and dynamics (**Fig. 4B**). Together with the results from the cysteine trapping analysis, this data suggests that both extracellular openings are a likely route for glycolipids exit, whereas the cytoplasmic lateral opening lined by TM2 and TM11 is more relevant for function as it may provide more room for glycolipids to enter the translocation pathway.

### The hydrophobic pocket is relevant for lipid transport

Inspection of the central cavity in the *in silico* analysis shows that similarly to what was observed in the outward-facing crystal structure of LtaA [26], the central cavity of the inward-facing state is amphipathic (**Fig. 5A**). The cavity displays a hydrophilic pocket, enclosed mainly by residues from the N-terminal domain (E32, R35, D68, W127 and W150), which we have previously shown to be relevant for recognition of the glycolipid headgroup [26], and a hydrophobic pocket, enclosed mainly by residues from the C-terminal domain (V234, L237, C263, L296, L300, L309, I316, and Y320) (**Fig. 5A**). The recent structure of the MFSD2A transporter trapped in an inward-facing conformation, displays a similar amphipathic central cavity (**Suppl. Fig. 6**) [21]. Computational docking of a glycolipid molecule to inward-facing LtaA suggests that the gentiobiosyl headgroup is preferentially accommodated in the hydrophilic pocket, whereas the diacylglycerol aliphatic tails are docked into the hydrophobic pocket (**Fig. 5B**).

**Figure 5.**
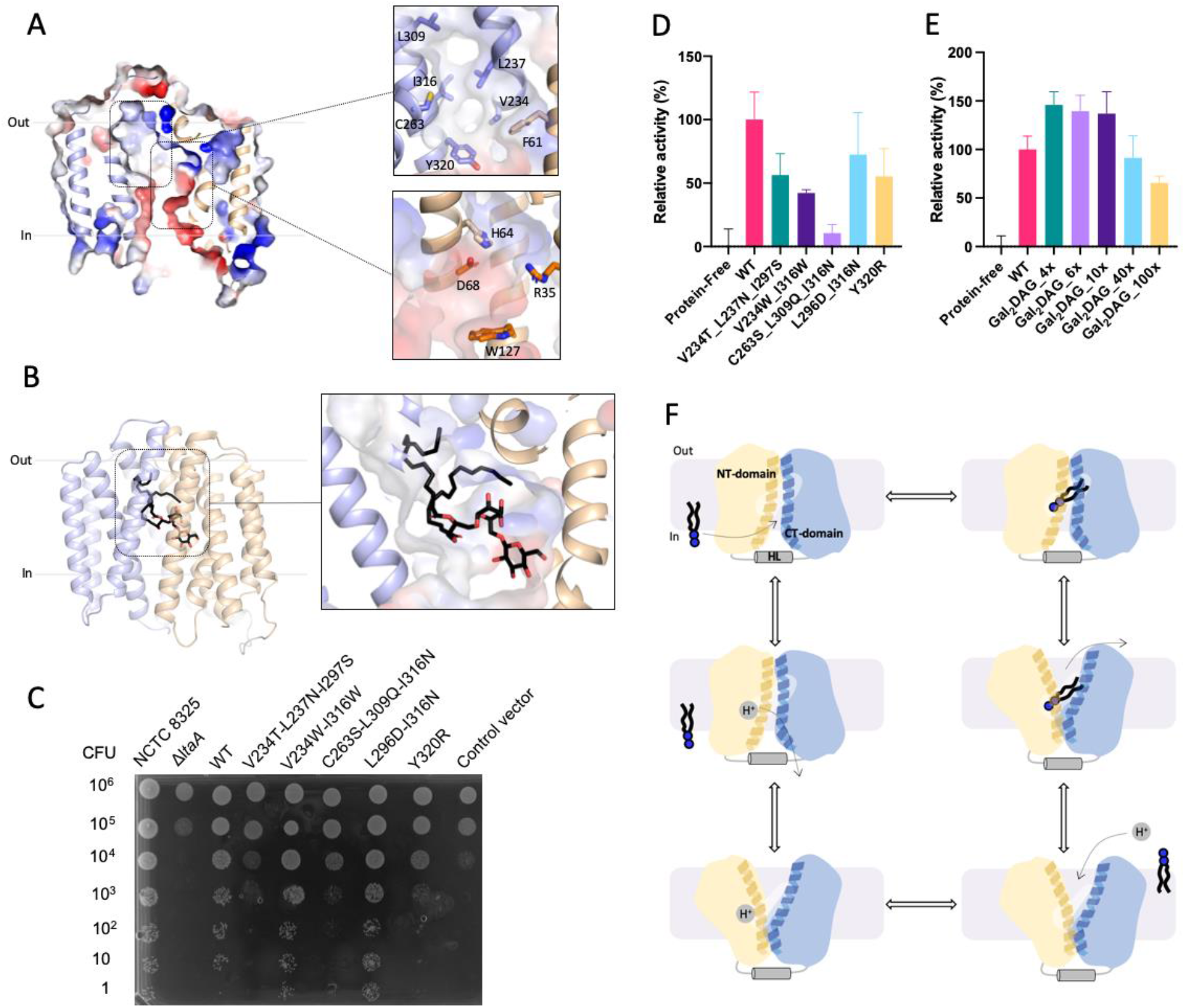
Hydrophilic and hydrophobic cavities participate in ‘trap-and-flip’ of lipids. **A**. Vacuum electrostatic surface representation of inward-facing model of LtaA. Residues forming the hydrophobic and hydrophilic pockets are shown. **B**. A model of a glycolipid molecule docked into the amphipathic cavity of LtaA. The lipid tail length corresponds to C16 chains. N-terminal and C-terminal domains are shown in light orange and light blue, respectively. **C**. *S. aureus* cell growth on LB agar plates containing 0.1 mM IPTG, buffered at pH 6.4. The *ΔltaA* mutant is complemented with pLOW vector carrying a *ltaA*-WT gene or the annotated point mutations; Control vector indicates the pLOW vector carrying an unrelated gene (dCas9). **D.** Mutagenesis analysis of the hydrophobic pocket. Relative flipping activity of LtaA-WT and variants. Error bars show s.d. of technical replicates, n≥3. **E**. Headgroup selectivity analysis. Relative flipping activity of LtaA in the presence of different concentrations of digalactosyl-diacylglycerol (Gal_2_DAG). Molar excess of Gal_2_DAG over Glc_2_-DAG-NBD is indicated. Error bars show s.d. of technical replicates, n≥3. **F**. Proposed mechanism of LtaA catalyzed glycolipid transport. Schematic of conformational states throughout LtaA transport cycle.

A striking feature of the central cavity observed in LtaA and MFSD2A [21], and to our knowledge, not observed in other MFS structures available to date, is the presence of the highly hydrophobic pocket at the C-terminal domains of these transporters (**Fig. 5A and Suppl. Fig. 6**). To test the importance of this pocket in LtaA, we have designed mutants that introduce polar residues, thus making it more hydrophilic. We then evaluated growth of *S. aureus* NCTC8325 *ΔltaA* cells complemented with ectopic copies of the *ltaA* gene carrying these mutations (**Fig. 5C and Suppl. Fig 7**). The variants V234T/L237N/I297S, C263S/L309Q/I316N, and Y320R display marked growth defects, whereas the mutant L296D/I316N do not affect growth. Each mutant was also purified and reconstituted into proteoliposomes, followed by determination of their flipping activity (**Fig. 5D**). In agreement with the results from *S. aureus* growth assays, the mutants V234T/L237N/I297S, C263S/L309Q/I316N, and Y320R display low relative activity compared to LtaA-WT (**Fig. 5D**), whereas L296D/I316N display the highest activity among all mutants. In contrast, introducing a mutation that scarcely increases the polarity of the cavity but that changes the size of residues V234 and I316, displayed low relative flipping activity compared to LtaA-WT, but does not affect growth of *S. aureus* NCTC8325 *ΔltaA* cells (**Fig. 5C,D and Suppl. Fig 7**). Taken together, these results support a fundamental role of the hydrophobic pocket in glycolipid transport. As suggested by docking analysis, it is likely that this pocket is involved in binding of the aliphatic tails of the glycolipid substrate. The striking hydrophobicity of the C-terminal TM helices 7, 8, and 10 in multiple MFS lipid transporters (**Suppl. Fig. 8**), and the involvement in coordination of the aliphatic chain of lysophospholipid as revealed by the structure of MFSD2A [21], suggest a shared mechanistic role of the hydrophobic pocket in lipid-tails binding in MFS lipid transporters.

#### The hydrophilic pocket dictates substrate specificity

So far, our results suggest that during transport, LtaA encloses the full glycolipid substrate in the amphipathic cavity. However, understanding the relevance of the individual parts of the substrate molecule, headgroup and aliphatic chains, is fundamental for future design of pharmacologically relevant molecules targeting this and other MFS lipid transporters. To gain insight into whether LtaA displays higher selectivity towards the headgroup than for the diacylglycerol moiety, we performed flipping assays with LtaA-WT co-reconstituted in proteoliposomes together with NBD-labeled Glc_2_-DAG (gentiobiosyl-diacylglycerol) and increasing concentrations of Gal_2_-DAG (digalactosyl-diacylglycerol) (**Fig. 5E**). Glucose and galactose differ only in the orientation of the -OH group at the C-4 position. Thus, we hypothesized that if the headgroup is more relevant for substrate recognition than the aliphatic chains, then transport of Glc_2_-DAG-NBD will be not be affected, since the difference between glucose and galactose would preclude Gal_2_-DAG from being a good competitor. On the other hand, if the diacylglycerol moiety is more relevant for substrate recognition, we expect Gal_2_-DAG to be a strong competitor, thus, resulting in marked decrease of Glc_2_-DAG-NBD transport. Our results show that even under a high excess of Gal_2_-DAG, there is no significant effect on Glc_2_-DAG-NBD transport (**Fig. 5E**). We have previously shown that gentiobiose (β-D-Glc-(1,6)-D-Glc), a disaccharide with the same composition and conformation as the glycolipid headgroup (Glc_2_-DAG), inhibits lipid transport [26]. Taken together, these results suggest that an intact headgroup is highly relevant for substrate binding and transport, and that even small changes to the headgroup abolishes recognition. Independent of the presence of the diacylglycerol moiety and its predicted binding to the hydrophobic pocket, the headgroup seems to dictate whether a glycolipid can be a substrate for LtaA or not.

## Discussion

Several transporters of the MFS superfamily have been structurally characterized in one or multiple conformational states [4, 8–17] However, except for the outward-facing structure of LtaA [26], solved by X-ray crystallography, and the inward-facing structure of MFSD2A [21], solved by single particle cryo-electron microscopy, there are no additional structures available of MFS lipid transporters. Despite the differences among their lipid substrates, the distinct composition of bacterial and eukaryotic membranes, and their opposite vectorial lipid transport directions, LtaA and MFSD2A share multiple architectural similarities, including a canonical MFS fold of 12 TM helices and an amphipathic central cavity with asymmetric distribution of hydrophobic and hydrophilic residues (**Suppl. Fig. 6**). A similar arrangement of central cavity residues has been predicted to be present in the bacterial lysophospholipid transporter LplT [18], and are likely to be part of the architecture of other MFS lipid transporters (**Suppl. Fig. 8**). These characteristics suggest a common mechanism of substrate recognition and translocation among these proteins. Indeed, LtaA and MFSD2A display strong selectivity towards the headgroup of their lipid substrates [6, 26]. In the case of MFSD2A the zwitterionic charge of the phosphatidylcholine headgroup is fundamental for ligand transport, whereas LtaA displays strong selectivity towards the gentiobiosyl disaccharide headgroup of the glycolipid. Furthermore, LtaA selects against an isomer of the disaccharide headgroup as shown by the poor competition displayed by digalactosyl-diacylglycerol in transport assays (**Fig. 5E**). In contrast, LplT has been shown to exhibit a more relaxed specificity towards the lipid headgroup, being able to transport lysophosphatidylethanolamine and lysophosphatidylglycerol lipids [18].

Although MFSD2A and LplT have been shown to strongly select for lysophospholipids, they display relaxed selectivity towards the length of the aliphatic chains [6, 18]. MFSD2A transports docosahexaenoic acid (DHA), an essential omega-3 fatty acid for brain growth and cognitive function, in the form of lysophosphatidylcholine, but can also transport other lipids with at least 14-carbons acyl chain [6]. It is noteworthy that *S. aureus* membranes are rich in diacylglycerols with chains length ranging from C_15_ to C_18_, with the most dominant lipid species having a C_17_:C_15_ composition [25]. This variability among diacylglycerols in *S. aureus*, and the measurable translocation of Glc_2_-DAG-NBD [26], which has a C_10_ acyl chain length and an NBD group linked to one of the diacylglycerol chains, suggest that LtaA displays similar relaxed specificity towards the length of the lipid part.

Our results strongly suggest that in contrast to mechanisms proposed for other lipid transporters, LtaA transports gentiobiosyl-diacylglycerol by a ‘trap-and-flip’ mechanism, which follows the classical alternating-access model of transport [51], with the entire glycolipid entering and leaving the central translocation pathway (**Fig. 5F**). Inward-facing LtaA binds a glycolipid molecule which enters through the lateral opening lined by TM2 and TM11, as suggested by the cross-linking analysis and MD simulations. This triggers a conformational change to an outward-facing state, in which the glycolipid is released into the membrane presumably through any of the two extracellular lateral openings. Protonation of residues in the hydrophilic pocket allows transition to the inward-facing conformation, followed by proton release to the cytoplasm.

Similar to the asymmetric opening of extracellular and cytoplasmic cavities described here for LtaA, MFSD2A exhibits a wider opening of its cytoplasmic lateral openings [21]. In this case, the extracellular opening is predicted to be narrow due to the constriction imposed by a disulfide bond at the extracellular side of the transporter [21]. Our results suggest that LtaA exhibit wide open extracellular lateral openings, whereas the cytoplasmic openings are narrower. The distinct constrictions of the cavities that recruit the lipid substrate, cytoplasmic cavity in LtaA and extracellular cavity in MFSD2A, might be part of a selectivity filter that confers substrate specificity.

In summary, our results provide insights into the molecular mechanism of glycolipid transport by LtaA and support a ‘trap-and-flip’ model where asymmetrically open lateral ‘gates’ and extracellular and cytoplasmic cavities play an essential role. Our data suggests that the highly selective hydrophilic pocket dictates substrate specificity, but that the hydrophobic pocket is fundamental for aliphatic chains transport. The mechanistic elements described here might be shared by other MFS lipid transporters and can be decisive for the design of drugs targeting these proteins.

## Methods

### LtaA expression and purification

The gene encoding *S. aureus* LtaA was cloned into a modified pET-19b vector (Novagen), with an N-terminal His_10_ affinity tag. LtaA WT and mutants were expressed in *E. coli* BL21 Gold (DE3) (Stratagene) cells. Cells were grown in Terrific Broth (TB) medium supplemented with 1% glucose (w/v) at 37°C. Overexpression was induced with 0.2 mM Isopropyl β-D-1-thiogalactopyranoside (IPTG) for 1h. All following steps were performed at 4 °C, unless different specified. Cells were harvested by centrifugation, resuspended in 50mM Tris-HCl, pH 8.0; 500mM NaCl; 5mM β-mercaptoethanol; 0.5mM PMSF and disrupted in a M-110L microfluidizer (Microfluidics) at 10000 psi chamber pressure. Membranes were pelleted by ultracentrifugation and solubilized in 50 mM Tris-HCl, pH 8.0; 200mM NaCl; 20mM Imidazole; 15% glycerol (v/v); 5mM β-mercaptoethanol; 1% Lauryl Maltose Neopentyl Glycol (w/v) (LMNG, Anatrace); 1% N-dodecyl-β-D-maltopyranoside (w/v) (DDM, Anatrace). After removing debri, the supernatant was loaded onto a pre-equilibrated NiNTA superflow affinity column (Qiagen). The column was washed with 50mM Tris-HCl, pH 8.0; 200mM NaCl; 50mM Imidazole; 10% glycerol (v/v); 5mM β-mercaptoethanol; 0.02% LMNG and 0.02% DDM and then further washed with the same buffer only containing 0.02% LMNG. Elution was performed in the same buffer containing 200mM Imidazole. Buffer exchange to buffer 10 mM Tris-HCl pH 8.0; 150mM NaCl; 0.02% LMNG was performed using PD-10 columns (GE Healthcare). Analytical size exclusion chromatography was performed on a Superdex 10/300 GL column (GE Healthcare) in buffer 10 mM Tris-HCl, pH 8.0; 150 mM NaCl; 0.02% LMNG. [52]

### Mutagenesis

LtaA mutants were generated using overlap Extension-PCR, followed by DpnI digestion for two hours at 37°C, and transformation into *E. coli* DH5α cells. The mutations were confirmed by DNA sequencing (Microsynth). All oligos used for mutagenesis are listed in **Suppl. table 2**.

### YpfP expression and purification

The gene encoding *S. aureus* YpfP was cloned into a modified pET-19b vector (Novagen) with an N-terminal His_10_ affinity tag. YpfP was overexpressed in BL-21 Gold (DE3) (Stratagene) cells. Cells were grown in TB medium supplemented with 1% glucose (w/v) at 37 °C until a cell density of OD_600_ = 3. Subsequently, cells were induced with 0.2 mM IPTG for 16h at 24 °C. Cells were harvested by centrifugation and resuspended in buffer A (50mM Tris-HCl pH 8.0; 200mM NaCl; 3% glycerol; 3mM β-mercaptoethanol) plus 0.5mM PMSF. Cells were disrupted using a tip sonication. After differential centrifugation, the supernatant containing YpfP was incubated with NiNTA resin and left stirring for 1h at 4 °C. Washing was performed with buffer A complemented with 50 mM imidazole pH 8.0, followed by elution with buffer A complemented with 200 mM imidazole pH8.0. YpfP was desalted in buffer 50 mM Tris-HCl pH 8.0; 200 mM NaCl; 10% glycerol using PD-10 columns (GE healthcare). If required YpfP was concentrated using a Vivaspin 20 30MWCO until 2.4 mg/ml, flash frozen in liquid nitrogen and stored at −80 °C until further use.

### Synthesis of NBD-glycolipid and glycolipid

Synthesis of glycolipd and nitrobenzoxadiazole (NBD)-labelled glycolipid was performed using a modification of the protocol described by Jorasch et al [53] and Kiriukhin et al [54]. A final concentration of 2mM UDP-Glucose (Sigma), 2mM NBD-decanoyl-2-decanoyl-sn-Glycerol (Cayman), and 1.2mg/ml purified YpfP were incubated together for 16h at 30°C. The reaction product was separated using thin-layer chromatography (TLC) with a silica gel matrix (Sigma) in a solvent mixture consisting of chloroform:methanol:water (65:25:4, vol/vol/vol). Silica containing the NBD-glycolipid was recovered from plates, and the NBD-glycolipid was extracted from the silica by incubation with a solvent mixture of chloroform:methanol (50:50, vol/vol), followed by drying of the anchor-LLD under argon atmosphere, and subsequently resuspension in 20 mM Tris-HCl pH 8.0; 150 mM NaCl. NBD-glycolipid was flash frozen in liquid nitrogen, and stored at −80°C until further use. Reaction products were previously characterized [26]. Non-labelled glycolipid was prepared similarly by incubation of 2mM UDP-Glucose, 2 mM 1,2-dimyristoyl-sn-glycerol (Avanti) and 1.2 mg/ml YpfP for 16h at 30°C.

### Formation of LtaA proteoliposomes

LtaA was reconstituted in unilamellar liposomes prepared by extrusion through polycarbonate filters (400 nm pore size) from a 3:1 (w/w) mixture of *E. coli* polar lipids and L-α-phosphatidylcholine (Avanti polar lipids) resuspended in 20 mM Tris-HCl pH 8.0; 150mM NaCl and 2mM β-mercaptoethanol. After saturation with DDM (Anatrace), liposomes were mixed with purified LtaA in a 50:1 (w/w) lipids/protein ratio. DDM was removed after incubation with BioBeads (BioRad). Proteoliposomes were centrifugated, washed and resuspended to a final concentration of 20mg/ml lipids; 7.8μM LtaA. The proteoliposomes were flash-frozen in liquid nitrogen and stored at −80°C until further use.

### *In vitro* flipping assay

Before performing flipping assays, proteoliposomes were thawed, their resuspension buffer was exchanged to 20 mM MES pH 6.5; 150 mM NaCl, and the product of the NBD-glycolipid synthesis reaction was incorporated by performing freeze/thaw cycles. Proteoliposomes and protein-free liposomes were diluted to a concentration of 2 mg/ml lipids followed by extrusion through polycarbonate filters (400 nm pore size). Proteoliposomes were immediately used for flipping assays. In case of competition assays with digalactosyldiacylglycerol (DGDG). DGDG powder (Avanti) was resuspended in 20 mM Tris-HCl; 150 mM NaCl and incorporated into proteoliposomes during freeze/thaw cycles together with the NBD-glycolipid. Flipping of NBD-glycolipid was assessed by determining the percentage of NBD-fluorescence that is quenched after addition of a 5 mM sodium dithionite (Sigma) after 200 seconds of starting fluorescence recording. 100 seconds before finishing data recording, 0.5% Triton X100 was added to permeabilize the liposomes, making all NBD-glycolipid molecules accessible to dithionite reduction. The fluorescence after Triton X100 addition was used for baseline calculations. Fluorescence was recorded at 20°C using a Jasco Fluorimeter. The excitation and emission wavelengths were 470 and 535 nm, respectively. For analysis the fluorescence intensity was normalized to F/F_max_. Relative flipping activities were calculated as follows: relative activity = 100 × ((F/F_max_)_i_ – (F/F_max_)_liposomes_) / ((F/F_max_)_wt_ – (F/F_max_)_liposomes_), where i corresponds to each respective treatment/mutants, liposomes corresponds to liposomes without protein, wt corresponds to wild type LtaA and F/Fmax values correspond to the normalized fluorescence values at the plateau after addition of sodium dithionite. Curves were plotted using GraphPad Prism 8. Time courses of the dithionite-induced fluorescence decay in liposomes were repeated at least 3 times for each individual experiment.

### Proton-transport assay

LtaA proteoliposomes and protein-free liposomes were thawed, and their internal buffer exchanged to 5 mM HEPES pH7.3; 100mM KCl. Glycolipid was incorporated during freeze/thaw cycles followed by extrusion through polycarbonate filters (400 nm pore size). After 90s of sonication, proteoliposomes and protein-free liposomes were diluted 25-fold in buffer containing 5 mM HEPES pH 7.3; 10 mM KCl; 90 mM NaCl; 0.5 μM 9-amino-6-chloro-2-methoxyacridine (ACMA). Fluorescence was recorded using a Jasco Fluorimeter with excitation and emission wavelengths of 410 and 480 nm respectively. When the fluorescence signal was stable, H^+^ influx was initiated by establishing a membrane potential by the addition of the potassium ionophore valinomycin (5 nM). Time courses of the proton-transport assay in proteoliposomes were repeated at least 3 times for each individual experiment. Crosslinking was performed before the measurement by addition of 2 mM CuCl_2_ to the proteoliposomes during the buffer exchange and incorporation of glycolipid steps. After 1h incubation at RT in the dark, CuCl_2_ was removed by centrifugation, and proteoliposomes were resuspended in buffer 5 mM HEPES, pH7.3; 100mM KCl.

### LtaA crosslinking and PEGylation

LtaA mutants incorporated into proteoliposomes were incubated with 2 mM CuCl_2_ or N,N’-1,2-phenylenedimaleimide (o-PDM) for 1h at RT in the dark. In case of non-crosslinked samples, proteoliposomes were incubated with a proportional volume of DMSO or buffer. Crosslinkers were removed by centrifugation and washing with buffer. To PEGylate free cysteines, LtaA mutants were incubated for 3h at RT in the presence of 0.5 mM mPEG5K-Maleimide (Sigma) and 0.5% SDS. Proteins were separated on 15% polyacrylamide gels and visualized with QuickBlue Protein stain (Lubio science).

### Sample preparation for LC-MS analysis

LtaA mutants were purified as described above, and concentrated to a concentration of 0.6 mg/ml. Purified LtaA was incubated for 1h at RT in the dark in the absence or presence of 2 mM o-PDM. Afterwards, 10 mM β-mercatoethanol was added to quench the crosslinker. 1-2 ug of either crosslinked or non-crosslinked LtaA protein were dissolved in 20 μl digestion buffer (0.02% of LMNG; 1 M urea; 0.1 M ammoniumbicarbonate; 10 mM tris(2-carboxyethyl) phosphine (TCEP); 15 mM chloroacetamide, pH = 8.5), reduced and alkylated for 1h at 37 °C. Proteins were digested by incubation with either sequencing-grade modified trypsin (1/50, w/w; Promega, Madison, Wisconsin), chymotrypsin sequencing grade (1/50, w/w, Sigma-Aldrich) or lys-C (1/100, w/w, Wako) overnight at 37°C. Then, the peptides were cleaned using iST cartridges (PreOmics, Munich) according to the manufacturer instructions. Samples were dried under vacuum and dissolved in 0.1 % formic acid solution at 0.5 pmol/ul. All samples were prepared in triplicates.

### Label-free targeted PRM-LC-MS analysis of cysteine-containing peptides

In a first step, parallel reaction-monitoring (PRM) assays [55] were generated for all the peptides of LtaA WT and the peptides of the 5 different LtaA cysteine mutants, for each protease. These peptides include the reference peptide for normalization, that is shared for all mutants. Therefore, the specific peptide sequences were loaded into Skyline (version 20.2.0.343 (https://brendanx-uw1.gs.washington.edu/labkey/project/home/software/Skyline/begin.view) and transitions were predicted using the integrated PROSIT algorithm for double and triple charged precursors. Then, protease and isoform specific isolation mass lists were exported and used to generate specific targeted LC-MS analyses. This analysis was carried as described previously [56]. Chromatographic separation of peptides was carried out using an EASY nano-LC 1000 system (Thermo Fisher Scientific), equipped with a heated RP-HPLC column (75 μm x 30 cm) packed in-house with 1.9 μm C18 resin (Reprosil-AQ Pur, Dr. Maisch). Aliquots of 1 pmol total peptides were analyzed per LC-MS/MS run using a linear gradient ranging from 95% solvent A (0.15% formic acid, 2% acetonitrile) and 5% solvent B (98% acetonitrile, 2% water, 0.15% formic acid) to 30% solvent B over 90 minutes at a flow rate of 200 nl/min. Mass spectrometry analysis was performed on a Q-Exactive plus mass spectrometer equipped with a nanoelectrospray ion source (both Thermo Fisher Scientific) using a hybrid DDA (top5)/PRM LC-MS analysis. In detail, each MS1 scan was followed by high-collision-dissociation (HCD) of the precursor masses of the imported isolation list and the 5 most abundant precursor ions with dynamic exclusion for 20 seconds. For each mutant and protease, a specific LC-MS method was generated. Total cycle time was approximately 1 second. For MS1, 3e6 ions were accumulated in the Orbitrap cell over a maximum time of 100 ms and scanned at a resolution of 70,000 FWHM (at 200 m/z). Targeted MS2 scans were acquired at a target setting of 3e6 ions, accumulation time of 100 ms and a resolution of 35,000 FWHM (at 200 m/z) and a mass isolation window to 0.4 Th. MS1 triggered MS2 scans were acquired at a target setting of 1e5 ions, a resolution of 17,500 FWHM (at 200 m/z) and a mass isolation window of 1.4 Th. Singly charged ions and ions with unassigned charge state were excluded from triggering MS2 events. The normalized collision energy was set to 27% and one microscan was acquired for each spectrum. The acquired raw-files were converted to mgf-file format using MSConvert (v 3.0, proteowizard) and searched using MASCOT (Matrix Science, Version: 2.4.1) against a decoy database containing normal and reverse sequences of the predicted SwissProt entries of *Staphylococcus aureus* (strain NCTC 8325 / PS 47, www.ebi.ac.uk, release date 2020/08/21). The 5 LtaA mutants and commonly observed contaminants (in total 6,574 sequences) were generated using the SequenceReverser tool from the MaxQuant software (Version 1.0.13.13). The search criteria were set as following: full tryptic specificity was required (cleavage after lysine or arginine residues); 3 missed cleavages were allowed; carbamidomethylation (C) was set as fixed modification and oxidation (M) as variable modification. The mass tolerance was set to 10 ppm for precursor ions and 0.02 Da for fragment ions. Then, Scaffold (version Scaffold_4.11.1, Proteome Software Inc., Portland, OR) was used to validate MS/MS based peptide and protein identifications. Peptide identifications were accepted if they could be established at a probability greater than 97.0% by the Scaffold Local FDR algorithm. Protein identifications were accepted if they could be established at a probability higher than 99.0% to achieve an FDR less than 1.0% and contained at least 2 identified peptides. Protein probabilities were assigned by the Protein Prophet algorithm [57]. Proteins that contained similar peptides and could not be differentiated based on MS/MS analysis alone were grouped to satisfy the principles of parsimony. Subsequently, all raw-files were imported into Skyline for protein/peptide quantification. To control for variation in sample amounts, all intensities were normalized against the 4 cysteine-free reference peptides. Only peptides that could be confidently identified by database searching were considered for quantification by PRM using the predicted transitions. Statistical analysis and ratio calculations to compare the relative abundance of the peptides between non-crosslinked and crosslinked peptides were performed in Excel. Histograms and P values were generated using Prism 9.

### Docking of glycolipid

A 1,2-dihexadecanoic-3-O-(β-D-glucopyranosyl-1→6-O-β-D-glucopyranosyl-sn-Glycerol molecule was docked to the LtaA inward-facing model with Autodock Vina [58]. The initial glycolipid coordinates were generated from 2D geometry in Phenix (eLBOW) [59]. The stereochemistry was corrected in Phenix (REEL)[59]. Docking was performed over a search space of 50×44×76 Å^3^ covering the central cavity.

### *S. aureus* phenotypic complementation assay

Generation of pLOW-ltaA and of *Staphylococcus aureus* NCTC8325 *ΔltaA* genotype was previously described [26]. pLOW vector was used for construction of *ltaA* complentary strains. Point mutations were generated by extension overlap PCR, and then with restriction-ligation cloning using SalI and NotI cloned into pLOW vector [60]. For cloning purposes *E. coli* IM08B was used[61]. The sequence of the resulting constructs was confirmed by DNA sequencing (Microsynth). After conformation of the correct constructs, pLOW vector carrying *ltaA* WT or point mutations were introduced into *S. aureus* NCTC8325 *ΔltaA* by electrophoresis with erythromycin selection (5μg/ml). *S. aureus* cells were grown in 3 ml of Luria-Bertani (LB) medium at 37 °C with 200 rpm until OD_600_ of 0.3. For complementary strains containing a pLOW vector, a final concentration of 5 μg/ml was added to the medium. For the serial dilutions, 5 μl of the original and its dilutions were spotted on LB agar plates buffered with sodium phosphate at pH 6.4 complemented with 0.1 mM IPTG. The plates were incubated overnight at 37 °C. Pictures were taken the next morning.

### Preparation of *S. aureus* membranes for LC-MS analysis

*S. aureus* cells were grown in 3 ml LB medium at 37 °C with 200 rpm until OD_600_ of 0.4. For complementary strains containing a pLOW vector, a final concentration of 5 μg/ml and 0.1 mM IPTG were added to the medium. After harvesting the cells were resuspended in 10 mM Tris pH8.0; 1 mM EDTA; 25 μg/ml lysostaphin, and incubated for 0.5h at 37 °C. Cells were further subjected to sonication, followed by collection of membranes by ultracentrifugation. The membranes were resuspended in 100 mM Tris-HCl; 5% SDS; 10 mM tris(2-carboxyethyl) phosphine (TCEP). Samples were sonicated for 10 minutes, followed by shaking for 1h at 37 °C with 500 rpm. To reduce and alkylate the disulfides a final concentration of 15 mM iodoacetamide was added, and the samples were incubated for 0.5h in the dark at room temperature. Samples were loaded on S-trap Micro Spin column (Protifi). After washing, on column peptide digestion was performed by addition of trypsin in 50 mM triethylammonium bicarbonate (TEAB) buffer, and incubation of 1h at 47 °C. Digested peptides were collected by passing 50 mM triethylammonium bicarbonate (TEAB) buffer, 0.2 % formic acid (w/v) in distilled water, and 0.2 % formic acid (w/v) in 50% acetonitrile (v/v) through the column and dried in a SpeedVac (Labconco). Dried peptides were re-suspended in 0.1% formic acid (w/v) and stored at −20 °C.

### Targeted PRM LC-MS analysis of LtaA WT and mutants

As a first step, PRM assays [55] for all possible peptides of LtaA with a length of 6 to 25 amino acids comprising double- and triple-charged precursor ions were created. Five peptides were identified to match the length and charge criteria, leading to ten PRM assays in total. These were used to identify LtaA membrane fractions of wild-type *S. aureus*. The setup of the μRPLC-MC system was previously described[56]. Mass spectrometry analysis was conducted using a Q-Exactive mass spectrometer with a nano-electrospray ion source (both Thermo Fisher Scientifice). Each MS1 scan was followed by high-collision-dissociation (HCD) of the ten LtaA precursor ions in PRM mode using a global isolation mass list. By applying strict identification criteria, three peptide ions of LtaA LTNYNTRPVK (2+ and 3+ ion) and MQDSSLNNYANHK (2+) were identified, and these were used for label-free PRM quantification. To control for protein variation between different samples, the total ion chromatography (only comprising peptide with two or more charges) was determined for each sample by label-free quantification using Progenesis QI (version 2.0, Waters) and then used to normalize the samples. The integrated peak areas of the three peptide ions that were quantified by PRM were summed up, and used for LtaA quantification.

### Modeling of inward-facing conformation

The inward-facing conformation was modelled under the assumption of inverted repeats [47]. Sequence alignments between the two repeats of each domain of LtaA were performed. We structurally aligned R1D1 (residues 16–105) with R2D1 (residues 109-189), and R1D2 (residues 220-302) with R2D2 (residues 309-393) using the structure alignment program TMalign resulting in two pairs of alignments. These two pairs of alignments were then used together to build up the final pair-wise alignment between the LtaA sequence and a template in which the LtaA sequence repeats were rearranged in the order R1D2-R1D1-R2D2-R2D1. The initial sequence alignment was then refined by removing gaps in the transmembrane regions and in the secondary structure elements. Further refinements were made to match the secondary structure as observed in the outward-open crystal structure. In particular, we aimed to maintain the helical regions in the template where possible, subject to the pseudo-symmetry between the two MFS transporter domains. We used this alignment and the X-ray crystal structure of LtaA (PDB entry 6S7V)[26] to construct the inward-facing model templates using Modeller 9v24. In 100 templates, the side chain packing of the models were re-built using SCWRL4 model. Next, we selected 7 models with the highest MODELLER score and the best MolProbity [62] profile for further analysis. Then, we repacked the side chains using SCWRL4.0[63] and as a last step the models were energetically minimized after placing them in the lipid bilayer using the Gromacs 2019.6 steepest descent algorithm for 5,000 steps [64]. To further validate the quality of the models, we assessed the stereochemistry. Evaluation of the model using MolProbity showed that the final minimized models have reasonable qualities (MolProbity score: 2.00-2.3, Ramachandran favored: 92.1-93.6%, and Ramachandran outliers: 0.8-1.90%) (**Suppl. Table 1)**.

### Molecular dynamics simulations of inward-facing conformation models

To study their dynamics, each of the optimized inward-facing models was placed in a heterogenous lipid bilayer (POPG (65%), diacylglycerol (20 %), cardiolipin (10 %), and gentiobiosyl-diacylglycerol (5 %)) and then solvated in TIP3P water with 150 mM NaCl. The all-atom CHARMM36m force field was used for lipids, ions, and protein [65–67]. All simulations were performed using GROMACS 2019.6 [64]. The starting systems were energy-minimized for 5,000 steepest descent steps and equilibrated first for 1 ns of MD simulations in a canonical (NVT) ensemble and then for 7.5 ns in an isothermal-isobaric (NPT) ensemble under periodic boundary conditions. The initial restrains on the positions of nonhydrogen protein atoms were 4,000 kJ·mol^-1^·nm^2^. During equilibration, these restraints were gradually released. Particle-mesh Ewald summation with cubic interpolation and a 0.12-nm grid spacing was used to treat long-range electrostatic interactions [68]. The time step was initially 1 fs, and was then increased to 2 fs during the NPT equilibration. The LINCS algorithm was used to fix all bond lengths [69].

Constant temperature was set with a Berendsen thermostat [70], combined with a coupling constant of 1.0 ps. A semi-isotropic Berendsen barostat [70] was used to maintain a pressure of 1 bar. During production runs, the Berendsen thermostat and barostat were replaced by a Nosé–Hoover thermostat [71] and a Parrinello–Rahman barostat [72] The unconstrained production trajectories were analyzed with Visual Molecular Dynamics (VMD) [73] and MDAnalysis package [74, 75]. A simulation of each inward-facing model was performed for 150 ns.

### Molecular dynamics simulations of outward-facing conformation

The outward-facing structure of LtaA (PDB ID 6S7V) was embedded in a lipid bilayer composed of POPG-DAG-CL-gentiobiosyl-diacylglycerol using CHARMM-GUI [76]. The system was then solvated in TIP3P water with 150 mM NaCl. The all-atom CHARMM36m force field was used for lipids, ions, and protein [65–67]. All simulations were performed using GROMACS 2019.6 [64]. Simulations were performed with similar protocols as described above for inward-facing models. The simulation of the outward-facing structure was performed for 960 ns.

## Acknowledgments

We thank Prof. Jan-Willem Veening for providing us the *S. aureus* NCTC8325, *S. aureus* NCTC8325 *ΔltaA, E. coli* IMO8B cells and the pLOW-vector. We thank Xiaochun Li Blatter for assistance in cell expression and membranes preparations. This work was supported by the Swiss National Science Foundation (SNSF) (PP00P3_198903 to C.P), the Helmut Horten Stiftung (HHS) (to C.P), and by the Max Planck Society and the German Research Foundation (SFB 807: Membrane Transport and Communication, to A.R.M and G.H). E.L. was funded by the Biozentrum International PhD Program and the HHS.

## Author Contributions

E.L. performed *in vitro* and *in vivo* biochemical characterization of LtaA and variants. A.R.M. performed computational analysis. C.P. supervised the biochemical analysis. G.H. supervised computational analysis. A.S. and E.L. performed mass spectrometry analysis. E.L., A.R.M, and C.P analyzed the computational, structural and functional data. C.P. conceived and directed the project. All authors contributed to manuscript writing and revision.

