## Supplementary_Information for "Evidence for a ‘trap-and-flip’ mechanism in a proton-dependent lipid transporter"

**This PDF file includes:**

Figs. S1 to S8
Tables S1 to S2

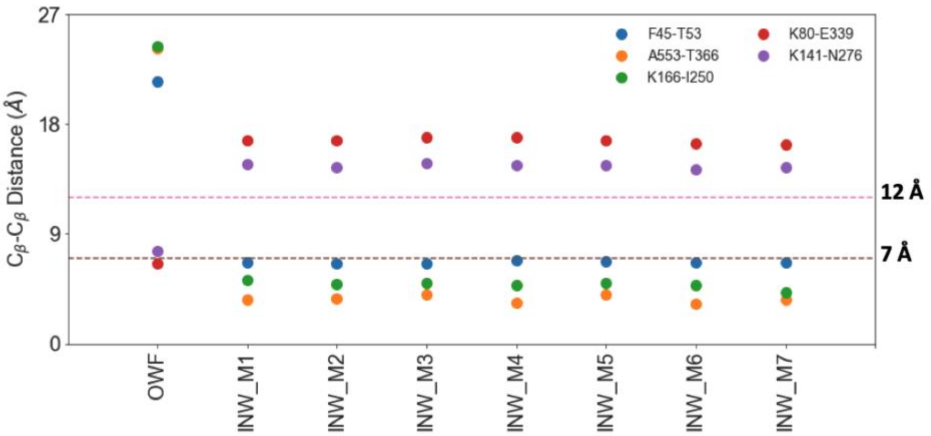


**Supplementary figure 1. Distances between residues selected for cross-linking in alternating conformations of LtaA.** Cβ-Cβ distances between residues selected for cysteine cross-linking in the outward-facing structure (OWF) and in the best inward-facing models (INW_M). Distance thresholds considered for selection of residues pairs are shown.

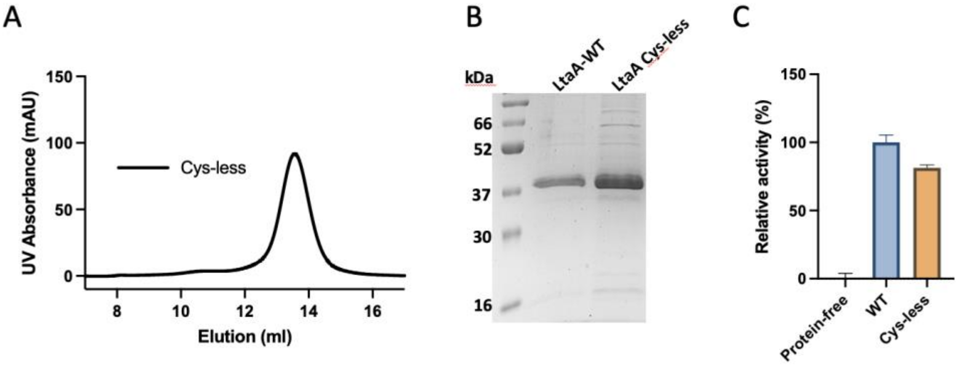


**Supplementary figure 2. Cysteine-less LtaA variant can be purified and displays activity similar to LtaA-WT. A**. Size Exclusion Chromatography profile of cysteine-less LtaA in a Superdex 200 Increase 10/300 GL column. **B**. SDS-PAGE of purified LtaA-WT and LtaA Cys-less. **C**. Relative flipping activity of LtaA Cys-less and LtaA-WT. Error bars show s.d. of technical replicates, n≥3.

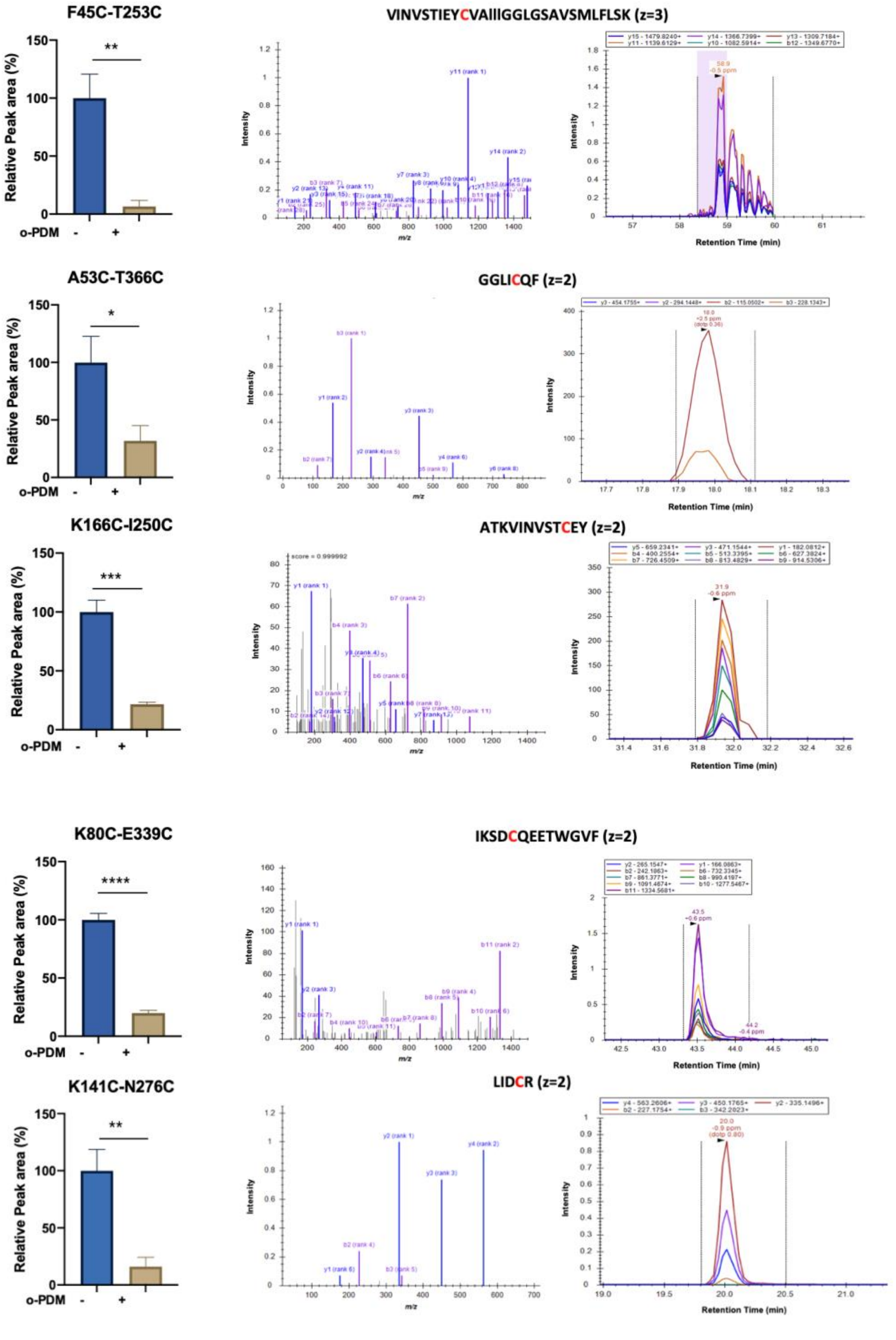


**Supplementary figure 3. LC-MS analysis of cysteine cross-linking showing that LtaA adopts inward- and outward-facing states.** Selected pairs of residues for cross-linking are shown in Fig. 2. *Left*: Relative abundance of cysteine containing peptides in absence (-) or presence (+) of N,N'-(o-phenylene)-dimaleimide (o-PDM). Error bars indicate s.d., n≥3. *: P≤0.05, **: P≤0.01, ***: P≤0.001, ****: P≤0.0001. *Center*: Assigned Collision-Induced Dissociation (CID) spectra obtained from cysteine containing peptides in absence of o-PDM. The position of the alkylated cysteine is indicated in red and the charge state (z) of the identified peptide ion is provided. *Right*: Elution profiles of the peptide fragments (transitions) identified used for peptide quantification by targeted LC-MS analysis.

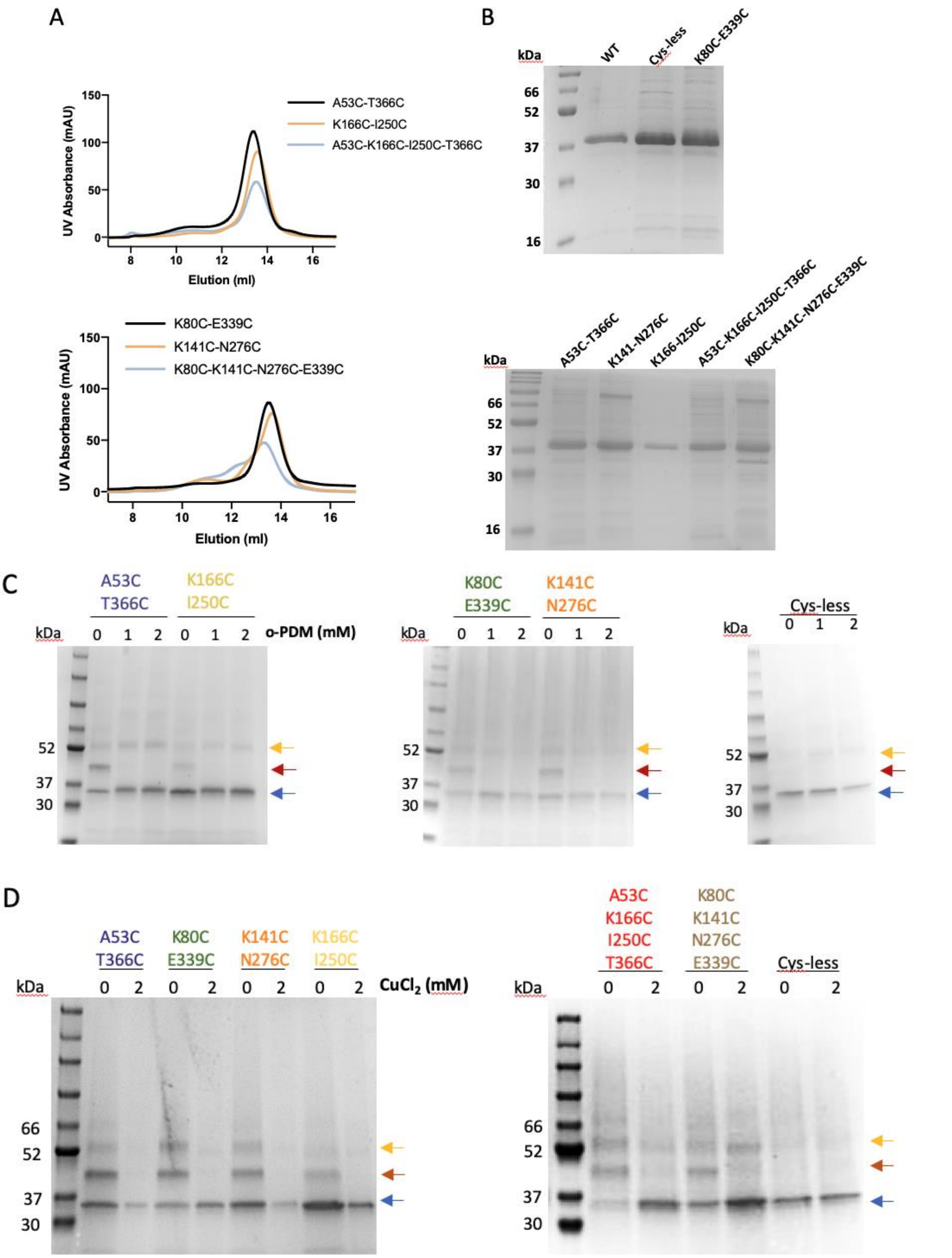


**Supplementary figure 4. Purifications and cross-linking in-gel analysis of LtaA variants. A**. Size Exclusion Chromatography profile of LtaA variants in a Superdex 200 Increase 10/300 GL column. **B**. SDS-PAGE of purified LtaA WT and variants. **C** and **D**. SDS-PAGE showing band shifts of samples treated with mPEG5K after cross-link with o-PDM or $CuCl_2$, respectively.

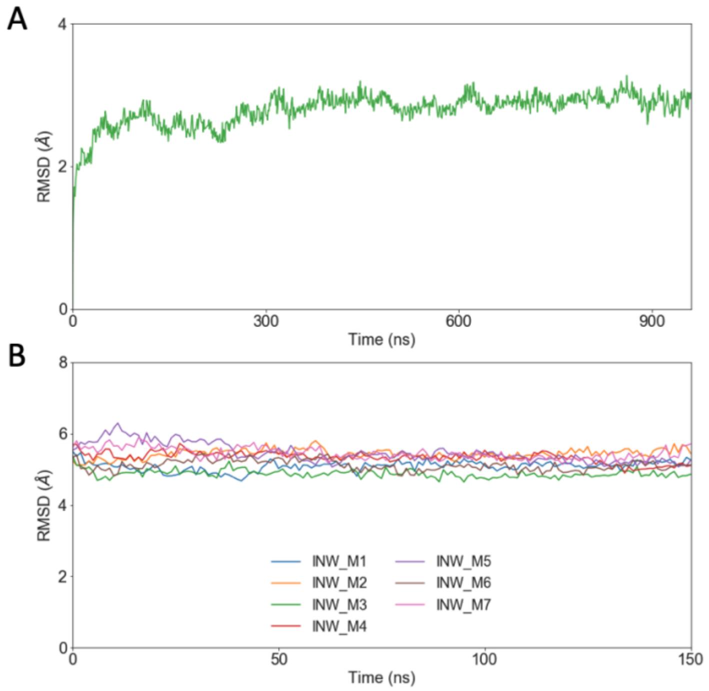


**Supplementary figure 5. RMSD analysis from molecular dynamics simulations of LtaA in a membrane.** Root-mean-square-deviation (RMSD) plots of Cα atoms from outward-facing LtaA (**A**) and inward-facing LtaA models (**B**) during MD simulations in a membrane composed of POPG (65%), diacylglycerol (20 %), cardiolipin (10 %), and gentiobiosyl-diacylglycerol (5 %).

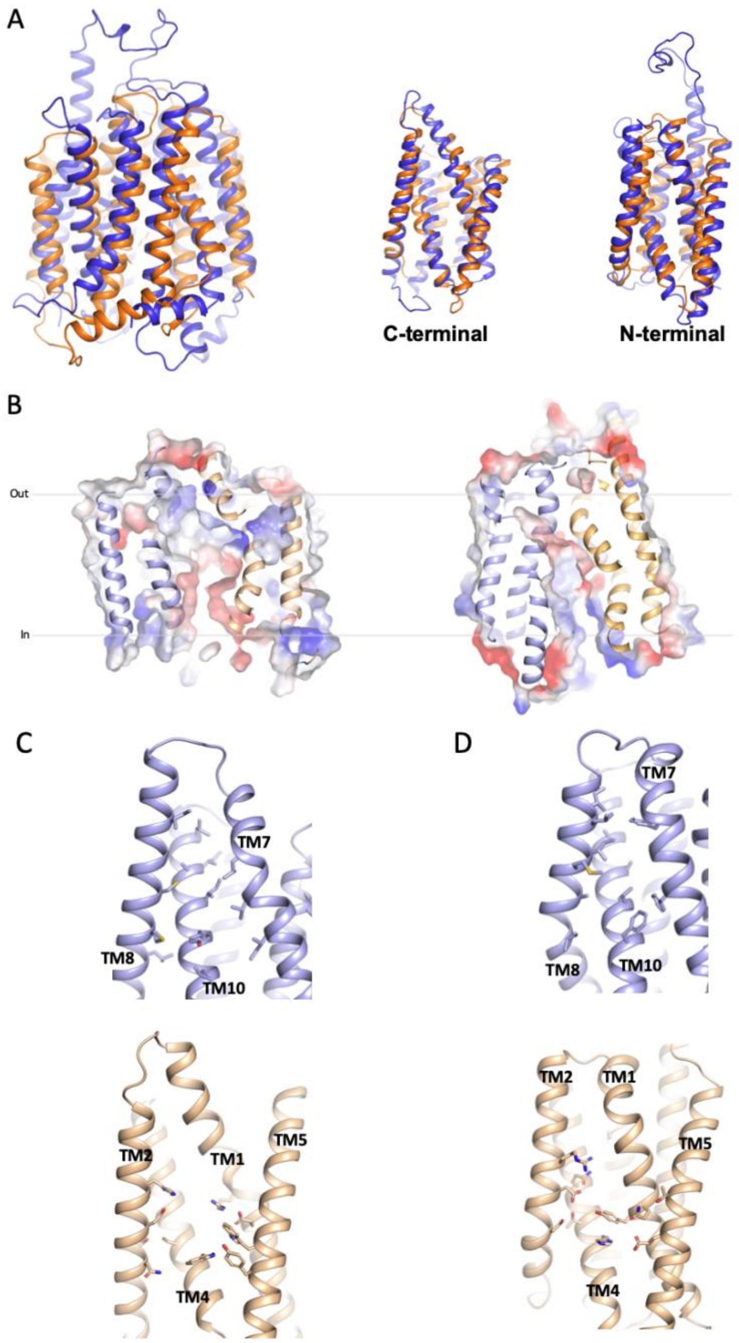


**Supplementary figure 6. Comparison of LtaA and MFSD2A fold and amphipathic cavity. A**. Superposition of LtaA (orange) and MFSD2A (blue) (r.m.s.d = 4.1Å), and individual N- and C-terminal domains (r.m.s.d= 2.6Å and 2.8Å, respectively). **B**. Vacuum electrostatic surface representation of inward-facing model of LtaA (*left*) and inward-facing MFSD2A (PDB ID 7MJS) (*right*). Residues participating in formation of N- and C-terminal hydrophilic and hydrophobic pockets in LtaA (**C**) and MFSD2A (**D**). N-terminal and C-terminal domains are shown in light orange and light blue, respectively.

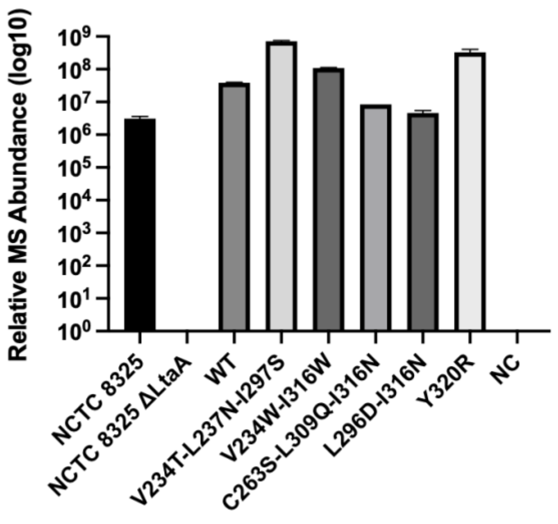


**Supplementary figure 7. LC-MS analysis of relative abundance of LtaA-WT and variants in *S. aureus* membranes.** The histogram shows relative abundances of LtaA WT and variants. Error bars show s.d. of technical replicates (n=3).

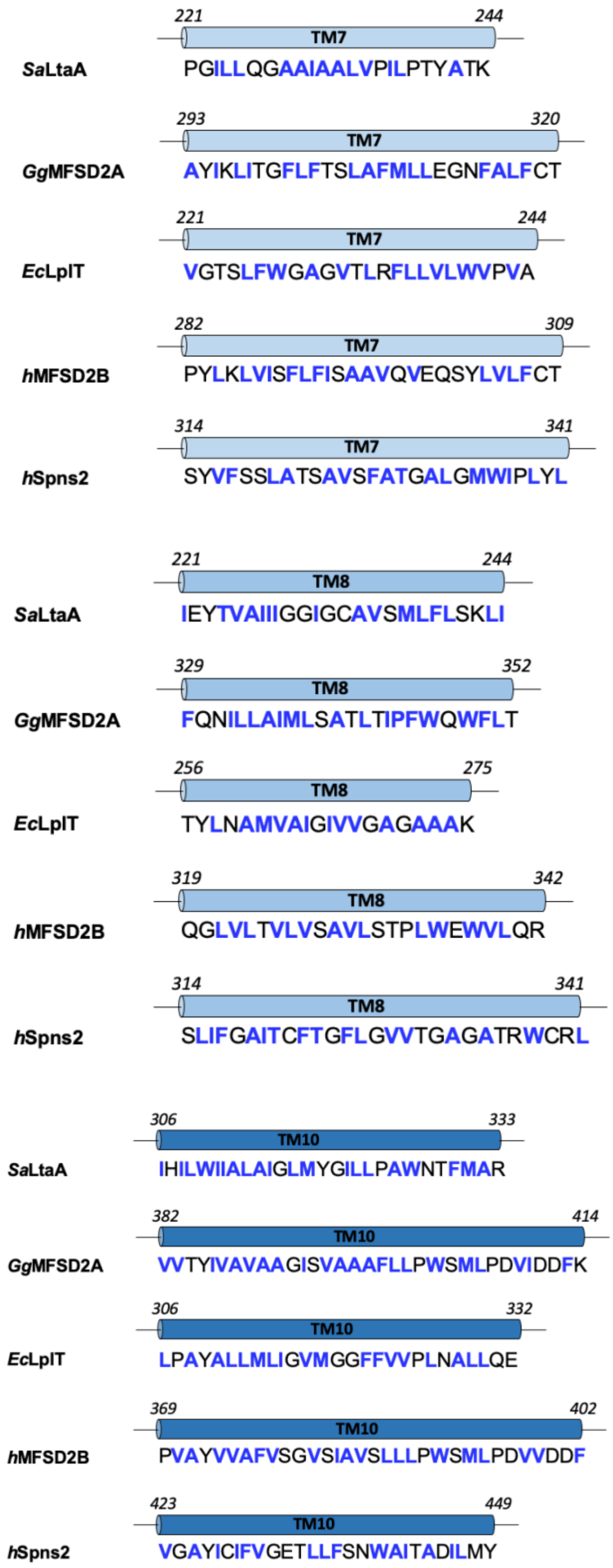


**Supplementary figure 8. C-terminal TM helices participating in formation of hydrophobic pockets in MFS lipid transporters.** Residues in TM7, 8, and 10 as seen in the structures of *S. aureus* LtaA (PDB ID 6S7V) and *Gallus gallus* MFSD2A (PDB ID 7MJS), and residues in predicted helical segments corresponding to the same TMs in *E. coli* LplT, human MFSD2B, and human Spns2 are shown. Hydrophobic residues are shown in blue.

**Supplementary Table 1.** Validation parameters of the top seven inward-facing models. *Initial*: homology models from Modeler: *SCWRL*; the refined model by SCWRL4; *mini*: energy minimization of the model in a bilayer using Gromacs.

| Model | Clashscore | | | MolProbity score | | | Ramachandran favoured (%) | | | Ramachandran outliers (%) | | |
|---|---|---|---|---|---|---|---|---|---|---|---|---|
| | Initial | SCWRL | mini | Initial | SCWRL | mini | Initial | SCWRL | mini | Initial | SCWRL | mini |
| 1 | 144.2 | 125.8 | 11.5 | 3.46 | 2.89 | 2.03 | 95.5 | 95.2 | 93.4 | 0.8 | 1.0 | 1.0 |
| 2 | 155.0 | 125.3 | 11.5 | 3.49 | 2.89 | 2.15 | 95.2 | 95.2 | 92.9 | 0.8 | 0.8 | 1.6 |
| 3 | 145.5 | 125.1 | 10.3 | 3.45 | 2.91 | 2.3 | 94.9 | 94.9 | 92.2 | 1.1 | 1.1 | 1.6 |
| 4 | 137.9 | 117.8 | 12.0 | 3.25 | 2.85 | 2.05 | 95.5 | 95.5 | 92.9 | 0.5 | 0.5 | 0.8 |
| 5 | 144.9 | 119.1 | 10.6 | 3.34 | 2.89 | 1.98 | 94.9 | 95.0 | 93.4 | 0.8 | 0.8 | 1.3 |
| 6 | 134.7 | 125.7 | 8.12 | 3.35 | 2.89 | 2.17 | 95.2 | 95.2 | 93.6 | 1.1 | 1.1 | 1.1 |
| 7 | 131.8 | 125.5 | 9.81 | 3.31 | 2.86 | 2.18 | 95.7 | 95.7 | 92.3 | 1.1 | 1.1 | 1.1 |

**Supplementary Table 2**. Oligo's used in this study. Mutagenesis is underlined

| Oligo name | 5'-3' sequence |
|---|---|
| Sa-SalI-FW | GAGAGTCGACGGTCATTCATCACAACC |
| Sa-V234T-L237N-FW | GGTGCCGCAATTGCTGCCCTAACCCCTATAAATCCAACATATGCTACTAAGGTTAT |
| Sa-V234T-L237N-RV | ATAACCTTAGTAGCATATGTTGGATTTATAGGGGTTAGGGCAGCAATTGCGGCACC |
| Sa-V234W-FW | GTGCCGCAATTGCTGCCCTATGGCCTATATTACCAACATATGCTAC |
| Sa-V234W-RV | GTAGCATATGTTGGTAATATAGGCCATAGGGCAGCAATTGCGGCAC |
| Sa-C263S-FW | CATTATTGGTGGTATCGGCTCTGCAGTTTCGATGCTATTT |
| Sa-C263S-RV | AAATAGCATCGAAACTGCAGAGCCGATACCACCAATAATG |
| Sa-L296D-FW | TCTAAGCGGATTTATTTTATACATGATAGATATTTTTACTCTATCTATGATTGTTAATATTCACA |
| Sa-L296D-RV | TGTGAATATTAACAATCATAGATAGAGTAAAAATATCTATCATGTATAAAATAAATCCGCTTAGA |
| Sa-I297S-FW | GATTTATTTTATACATGATATTAAGTTTTACTCTATCTATGATTGTTAATATTC |
| Sa-I297S-RV | GAATATTAACAATCATAGATAGAGTAAAACTTAATATCATGTATAAAATAAATC |
| Sa-L309Q-I316N-FW | CTATGATTGTTAATATTCACATCCAATGGATTATCGCTTTAGCTAATGGTCTAATGTATGGCATCTT ATTAC |
| Sa-L309Q-I316N-RV | GTAATAAGATGCCATACATTAGACCATTAGCTAAAGCGATAATCCATTGGATGTGAATATTAACA ATCATAG |
| Sa-I316N-FW | CTTGTGGATTATCGCTTTAGCTAATGGTCTAATGTATGGCATCTTATTAC |
| Sa-I316N-RV | GTAATAAGATGCCATACATTAGACCATTAGCTAAAGCGATAATCCACAAG |
| Sa-I316W-FW | CTTGTGGATTATCGCTTTAGCTTGGGGTCTAATGTATGGCATCTTATTAC |
| Sa-I316W-RV | GTAATAAGATGCCATACATTAGACCCCAAGCTAAAGCGATAATCCACAAG |
| Sa-Y320R-FW | TTAGCTATCGGTCTAATGCGTGGCATCTTATTACCAGCATG |
| Sa-Y320R-RV | CATGCTGGTAATAAGATGCCACGCATTAGACCGATAGCTAA |
| LtaA-NotI-RV | GAGAGCGGCCGCAATAGTATTGTTAATCGTAGTATGTTTGAATTAATAAGA |
| F45C-FW | GAGCTACATTAACTGTCTGCCCACCG |
| F45C-RV | CGGTGGGCAGACAGTTAATGTAGCTC |
| A53C-FW | CCGTGACATCGATTTGTGTCGCTATTACATC |

| | |
|---|---|
| A53C-RV | GATGTAATAGCGACACAAATCGATGTCACGG |
| K80C-FW | CGTTATCGGCTTCTTATTATGTAAGTTTGGAACTAAGATC |
| K80C-RV | GATCTTAGTTCCAAACTTACATAATAAGAAGCCGATAACG |
| K141C-FW | GAGGATAAACGCGGCTGTCAAATGGGCTACGTG |
| K141C-RV | CACGTAGCCCATTTGACAGCCGCGTTTATCCTC |
| K166C-FW | CTTTATGAATTTGCTGATCTGTGTTCATCCTACTCGCTTCG |
| K166C-RV | CGAAGCGAGTAGGATGAACACAGATCAGCAAATTCATAAAG |
| I250C-FW | CAAAAGTCATTAATGTATCAACGTGTGAATATACCGTGGCTATTATC |
| I250C-RV | GATAATAGCCACGGTATATTCACACGTTGATACATTAATGACTTTTG |
| T253C-FW | GTATCAACGATTGAATATTGTGTGGCTATTATCATCGGCGG |
| T253C-RV | CCGCCGATGATAATAGCCACACAATATTCAATCGTTGATAC |
| C263S-FW | CGGCGGCATTGGATCGGCCGTTAGTATGTTG |
| C263S-RV | CAACATACTAACGGCCGATCCAATGCCGCCG |
| N276C-FW | CTTAGTAAGTTGATCGACTGCCGTTCGCGTAACTTTATGTAC |
| N276C-RV | GTACATAAAGTTACGCGAACGGCAGTCGATCAACTTACTAAG |
| E339C-FW | GCTTCATCAAGAGCGACTGTCAAGAGGAGACCTGG |
| E339C-RV | CCAGGTCTCCTCTTGACAGTCGCTCTTGATGAAGC |
| T366C-FW | GTTTGGCGGATTGATTTGCCAGTTCACCAAC |
| T366C-RV | GTTGGTGAACTGGCAAATCAATCCGCCAAAC |
| LtaA-V234T-L237N-FW | GATTGCGGCTTTGACTCCTATCAACCCTACGTATGCCACA |
| LtaA-V234T-L237N-RV | TGTGGCATACGTAGGGTTGATAGGAGTCAAAGCCGCAATC |
| LtaA-V234W-FW | GATTGCGGCTTTGTGGCCTATCCTGCCTAC |
| LtaA-V234W-RV | GTAGGCAGGATAGGCCACAAAGCCGCAATC |
| LtaA-C263S-FW | GCGGCATTGGATCGGCCGTTAGTATG |
| LtaA-C263S-RV | CATACTAACGGCCGATCCAATGCCGC |
| LtaA-L296D-FW | GGTTTTATTCTGTATATGATTGATATCTTCACGTTGTCTATG |
| LtaA-L296D-RV | CATAGACAACGTGAAGATATCAATCATATACAGAATAAAACC |
| LtaA-I297S-FW | TTATTCTGTATATGATTCTTAGTTTCACGTTGTCTATGATTGTC |
| LtaA-I297S-RV | GACAATCATAGACAACGTGAAACTAAGAATCATATACAGAATAA |
| LtaA-L309Q-FW | GTCAACATCCATATCCAATGGATTATTGCCTTGG |
| LtaA-L309Q-RV | CCAAGGCAATAATCCATTGGATATGGATGTTGAC |
| LtaA-I316N-FW | TGCCTTGGCGAACGGCCTGATG |
| LtaA-I316N-RV | CATCAGGCCGTTCGCCAAGGCA |
| LtaA-I316W-FW | ATTATTGCCTTGGCGTGGGGCCTGATGTAT |
| LtaA-I316W-RV | ATACATCAGGCCCCACGCCAAGGCAATAAT |
| LtaA-Y320R-FW | GATCGGCCTGATGCGTGGGATTCTGTTGCCA |
| LtaA-Y320R-RV | GATCGGCCTGATGCGTGGGATTCTGTTGCCA |
